# Novel genetic traits in *Xanthomonas* unveiled by the complete genome sequencing of three clade-1 xanthomonads

**DOI:** 10.1101/2022.10.10.511549

**Authors:** Chloé Peduzzi, Angeliki Sagia, Daiva Burokienė, Ildikó Katalin Nagy, Marion Fischer-Le Saux, Perrine Portier, Alexis Dereeper, Sébastien Cunnac, Veronica Roman-Reyna, Jonathan M. Jacobs, Claude Bragard, Ralf Koebnik

**Affiliations:** Earth & Life Institute, UCLouvain, Louvain-la-Neuve, Belgium; Plant Health Institute of Montpellier (PHIM), University of Montpellier, Cirad, INRAE, Institut Agro, IRD, Montpellier, France; Nature Research Centre, Institute of Botany, Laboratory of Plant Pathology, Vilnius, Lithuania; Enviroinvest Corp., Pécs, Hungary; Univ Angers, Institut Agro, INRAE, IRHS, SFR QUASAV, CIRM-CFBP, F-49000 Angers, France; Department of Plant Pathology, The Ohio State University, Columbus, OH 43210, U.S.A.; Infectious Disease Institute, The Ohio State University, Columbus, OH 43210, U.S.A.

**Keywords:** Taxonomy, Evolution, Coronatine, Stomata, Lateral flagellum, Banana, Common bean, New Zealand flax

## Abstract

Evolutionary early-branching xanthomonads, also referred to as clade-1 xanthomonads, include major plant pathogens, most of which colonize monocotyledonous plants. Seven species have been validly described, among them the two sugarcane pathogens *Xanthomonas albilineans* and *Xanthomonas sacchari*, and *Xanthomonas translucens*, which infects small grain cereals, diverse grasses, but also asparagus and pistachio trees. Single-gene sequencing and genomic approaches indicated that this clade likely contains more, yet undescribed species. In this study, we sequenced representative strains of three novel species using long-read sequencing technology. *Xanthomonas* sp. pv. *phormiicola* strain CFBP 8444 causes bacterial streak on New Zealand flax, another monocotyledonous plant. *Xanthomonas* sp. strain CFBP 8443 has been isolated from common bean and *Xanthomonas* sp. strain CFBP 8445 originated from banana. Complete assemblies of the chromosomes confirmed their unique taxonomic position within clade 1 of *Xanthomonas*. Genome mining revealed novel genetic traits, hitherto undescribed in other members of the *Xanthomonas* genus. In strain CFBP 8444, we identified genes related to the synthesis of coronatine-like compounds, a phytotoxin produced by several pseudomonads, which raises interesting questions about the evolution and pathogenicity of this pathogen. In addition, strain CFBP 8444 was found to encode a second, atypical flagellar gene cluster in addition to the canonical flagellar gene cluster. Overall, this research represents an important step toward better understanding the evolutionary history and biology of early-branching xanthomonads.

---

The Gram-negative bacterial genus *Xanthomonas* constitutes a large group of mostly plant-associated bacteria, collectively causing diseases on hundreds of host plants, including economically important crops and ornamental plants. Thanks to increased genome sequencing efforts, the number of *Xanthomonas* genome sequences has increased tremendously in recent years, including strains of scientifically and/or economically relevant species and pathovars (Mansfield et al. 2012). But also understudied non-model organisms have been systematically sequenced, thus correcting taxonomically mis-assigned strains and leading to description of novel species (Vicente et al. 2017; López et al. 2018; Martins et al. 2020; Bansal et al. 2021; Dia et al. 2021; Mafakheri et al. 2022).

Currently, the List of Prokaryotic names with Standing in Nomenclature (LPSN; https://www.bacterio.net, assessed on 8 September 2022) includes 34 validly described *Xanthomonas* species (Parte et al. 2020). Previous work based on partial sequencing of the *gyrB* gene from poorly characterized pathovars and from unidentified xanthomonads, however, suggested the existence of additional species, which were at that time described as species-level clade strains, SLC 1 to SLC 7 (Parkinson et al. 2009). Strains from SLC 1 and 4 have been sequenced since then, corresponding to the proposed species *‘Xanthomonas cannabis’* and *‘Xanthomonas badrii’*, respectively (Zain & Roberts 1977; Jacobs et al. 2015).

Xanthomonads are classified into two major groups based on sequence comparisons (Hauben et al. 1997; Gonçalves & Rosato 2002; Parkinson et al. 2007; Ferreira-Tonin et al. 2012). SLC 1 to 4 belong to clade 2, while SLC 5 to 7 belong to clade 1, also known as the clade of early-branching species (Parkinson et al. 2007; 2009). Most of the clade-1 strains have been isolated from monocots, such as asparagus, banana, sugarcane, hyacinths, forage grasses, and small-grain cereals (Jacques et al. 2016). But there are a few exceptions: *Xanthomonas theicola* causes canker on tea plants, *Xanthomonas bonasiae* and *Xanthomonas youngii* were found on canker-symptomatic ornamental plants, and strains of *Xanthomonas translucens* have also been isolated from pistachio trees (Giblot-Ducray et al. 2009; Koebnik et al. 2021; Mafakheri et al. 2022). SLC-5 strain NCPPB 2654 (= CFBP 8443) was isolated from bean (*Phaseolus vulgaris*), the SLC-6 strain NCPPB 2983 (= CFBP 8444, LMG 702) originates from New Zealand flax (*Phormium tenax*), and seven SLC-7 strains had been collected from monocots like banana (*Musa* × *paradisiaca*), ginger (*Zingiber officinale*) or sugarcane (*Saccharum officinarum*), but also from cotton (*Gossypium* sp.) (Parkinson et al. 2009).

The SLC-6 bacterium *X. campestris* pv. *phormiicola* (Takimoto 1933) Dye 1978 was described as a bacterial pathogen of New Zealand flax (*Phormium tenax* Forst.), causing water-soaked lesions with chlorosis (Takimoto, 1933). The same bacterium was also found to induce hypertrophy on potato tuber tissues at the inoculated site (Tamura et al. 1987). The hypertrophic active substance isolated from the bacterial culture filtrate produced visible chlorotic lesions on the leaves of New Zealand flax, ryegrass and tobacco plant (Tamura et al. 1992). Biochemical and biophysical parameters (retardation factor on silica gel thin-layer chromatography and 1H nuclear magnetic resonance [NMR] spectrum) of the purified toxin coincided with those of coronatine, and it was concluded that the toxin produced by *X. campestris* pv. *phormiicola* is coronatine (Tamura et al. 1992). In a parallel study, liquid cultures of *X. campestris* pv. *phormiicola* were found to contain two analogues of coronatine lacking the cyclopropane ring structure, but no trace of coronatine (Mitchell 1991). Using NMR, mass spectroscopy and gas chromatography of hydrolysis products, these two compounds were identified as N-coronafacoyl-l-valine and N-coronafacoyl-l-isoleucine (Mitchell 1991). These were the first two reports of coronatine-like phytotoxins outside of the genus *Pseudomonas*.

To obtain insight into the genetics of coronatine production in *Xanthomonas* and to expand genomic information on clade-1 xanthomonads, we determined the complete genome sequences of the pathotype strain CFBP 8444^PT^ (ICMP 4294^PT^, LMG 702^PT^, NCPPB 2983^PT^), and of the specieslevel clade strains CFBP 8443 (NCPPB 2654) and CFBP 8445 (NCPPB 1131).

## Genome sequencing and assembly

From lyophilized stocks available at CIRM-CFBP (https://cirm-cfbp.fr/), single colonies of the three strains were isolated and grown on peptone-sucrose (10 g/liter peptone, 10 g/liter sucrose, 1 g/liter glutamic acid) agar media. Genomic DNA was extracted using the Genomic DNA buffer set and Genomic-tips following the manufacturer’s instructions (Qiagen, Valencia, CA, U.S.A.).

Multiplex library preparation in pools of eight strains, including strains CFBP 8443, CFBP 8444, and CFBP 8445, and simultaneous sequencing of eight strains on one SMRTCell was conducted with the PacBio Sequel technology at the GENTYANE genotyping platform (INRAE Clermont-Ferrand, France).

Sequence reads were *de novo* assembled with Flye 2.7 using the “--plasmids --iterations 2” parameters (Kolmogorov et al. 2019), yielding a single chromosome for each of the three strains. Genome sequences were functionally annotated using the NCBI Prokaryotic Genome Annotation Pipeline (PGAP version 6.2) (Li et al. 2021). The clade-1 xanthomonads genome project is available at NCBI as BioProject PRJNA865934. Annotation details are given in Table 1.

**Table 1.** BioSamples, GenBank accession numbers and annotation details for the clade-1 xanthomonads genome BioProject PRJNA865934.

| Strain | CFBP 8443 | CFBP 8444 | CFBP 8445 |
| --- | --- | --- | --- |
| Species-level clade | SLC 5 | SLC 6 | SLC 7 |
| NCBI BioSample | SAMN30147989 | SAMN30148166 | SAMN30161540 |
| NCBI GenBank accession number<br>(chromosomes) | CP102592 | CP102593 | CP102594 |
| Coverage | 200 x | 244 x | 208 x |
| Size (bp) | 5,238,480 | 5,314,063 | 4,869,984 |
| G+C content (%) | 68.96 | 68.64 | 69.25 |
| Predicted genes | 4,368 | 4,506 | 4,192 |
| Predicted CDS | 4,303 | 4,441 | 4,122 |
| Predicted RNA genes | 65 | 65 | 70 |
| Predicted pseudogenes | 44 | 72 | 30 |

## Phylogenetic relationships

To clarify the phylogenetic relationships within clade 1, including the three newly sequenced SLC strains, pairwise average nucleotide identities (ANI) were calculated on a Galaxy-implemented version of FastANI (Jain et al. 2018). This analysis included all publicly available genome sequences from clade 1, except for *X. translucens*, where only eight pathotype strains were chosen as representatives of the genetic diversity of this species (Goettelmann et al. 2022). The evolutionary history was inferred using the UPGMA method (Sneath & Sokal 1973), as implemented in MEGA11 (Stecher et al. 2020; Tamura et al. 2021). This analysis involved 75 genome sequences and confirmed the taxonomic position of the three SLC strains within clade 1 (Figure 1; Supplemental Table 1).

**Figure 1.**
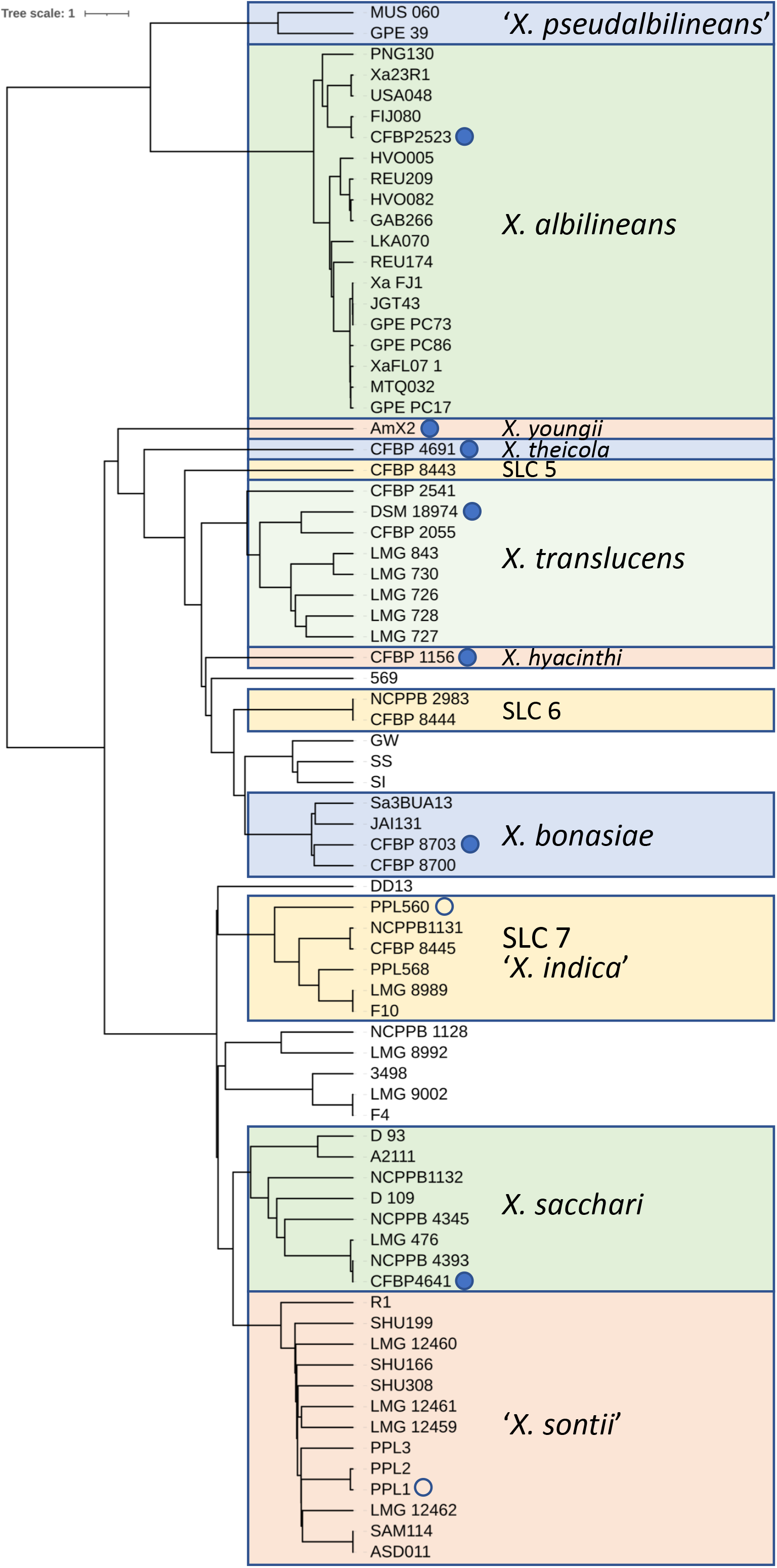
Phylogenetic tree of clade-1 xanthomonads. The evolutionary distances between genome sequences were computed using the FastANI method (Jain et al. 2018) and are in units of percent sequence divergence. The evolutionary history was inferred using the UPGMA method (Sneath & Sokal 1973). The tree is drawn to scale, with branch lengths in the same units as those of the evolutionary distances used to infer the phylogenetic tree. This analysis involved 75 genome sequences. Evolutionary analyses were conducted in MEGA11 (Stecher et al. 2020; Tamura et al. 2021), and the tree layout was enhanced for better visualization using the iTOL online tool (Letunic & Bork 2019). Names of bacterial strains are indicated next to the branch endpoints. Strains belonging to the same species are boxed with the name of the species given within the colored box. Type strains are labeled with a blue dot, and proposed type strains are labeled with a blue circle. Used genome sequences are listed in Supplemental Table 1.

Using a 95% cut-off value as an ANI species delimiter (Goris et al. 2007; Richter & Rosselló-Móra 2009), SLC-5 strain CFBP 8443 was found to be the only representative for this novel species. The SLC-6 strain CFBP 8444 and its clone NCPPB 2983 in the UK strain collection represent a sister species of *X. bonasiae* (Mafakheri et al. 2022). Three nearby strains in the dendrogram, GW, SI, and SS, which were isolated as candidate biocontrol agents from the microbiome of perennial ryegrass (*Lolium perenne*), were found to be phylogenetically close to the species *X. bonasiae* (Li et al. 2020). However, using digital DNA-DNA hybridization (dDDH) as a species delimiter suggests that strains GW, SI, and SS form another sister species of *X. bonasiae* (Supplemental Table 2) (Meier-Kolthoff et al. 2013, 2022). We therefore consider strain CFBP 8444 as the representative of a novel species, corresponding to SLC 6.

The SLC-7 strain CFBP 8445 is a clone of NCPPB 1131, for which a draft genome sequence was published previously (Studholme et al. 2011). These strains clustered with another genetically similar strain, LMG 8989, which was isolated from *Citrus* plants (Bansal et al. 2020), in contrast to CFBP 8445 and NCPPB 1131, which were isolated from banana. Recently, non-pathogenic strains were isolated from healthy rice seeds. Genome sequencing revealed that two of the strains, PPL560 and PPL568, belong to a novel species, for which the name *‘Xanthomonas indica’* was proposed (Rana et al. 2022). ANI and dDDH analyses demonstrate that these two strains belong to the same species-level clade as the four former strains and thus SLC 7 could be renamed ‘*X. indica’* (Figure 1, Supplemental Tables 1 and 2). Hence, this species-level clade comprises bacteria that can colonize both monocots and dicots. Whether such promiscuity also holds true for SLC 5 and SLC 6 strains awaits the characterization of additional isolates.

**Figure 2.**
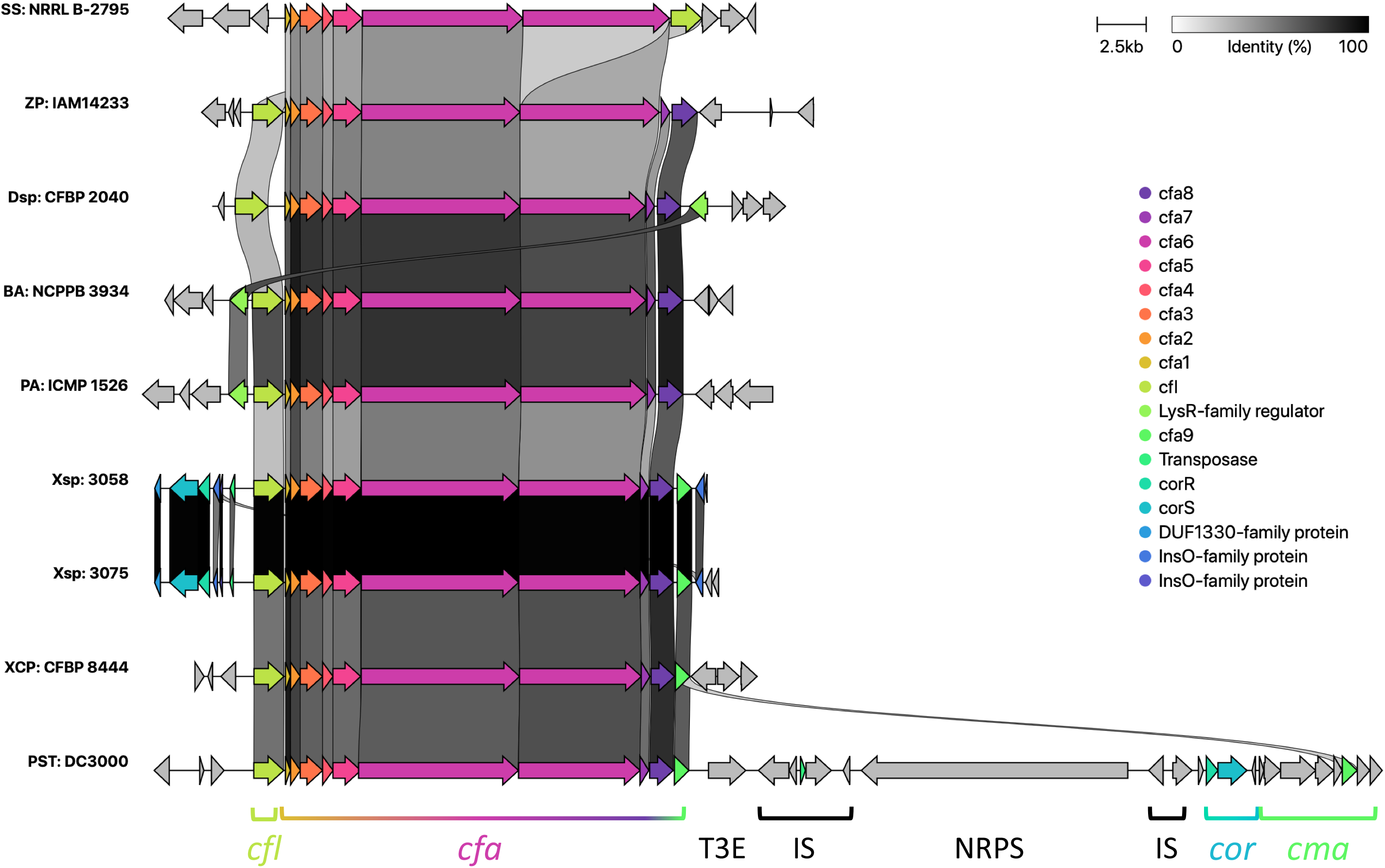
Comparison of bacterial coronatine biosynthesis gene clusters. Corresponding regions, extracted from GenBank files, were compared and visualized with clinker (Gilchrist & Chooi 2021). The following strains and genomes were analyzed (from top to bottom): *Streptomyces scabiei* strain NRRL B-2795 (JAGJBZ010000001), *Zymobacter palmae* strain IAM14233 (AP018933), *Dickeya* sp. strain CFBP 2040 (JAAVXH010000024), *Brenneria alni* strain NCPPB 3934 (MJLZ01000030), *Pectobacterium atrosepticum* strain ICMP 1526 (ALIV01000029), *Xanthomonas* sp. strain 3058 (JACHOF010000019), *Xanthomonas* sp. strain 3075 (JACIFI010000014), *Xanthomonas campestris* pv. *phormiicola* strain CFBP 8444 (CP102593), and *Pseudomonas syringae* pv. *tomato* strain DC3000 (AE016853). Genes and operons in strain DC3000 are labelled at the bottom. T3E = type 3 effector; NRPS = non-ribosomal polyketide synthase; IS = insertion sequence element.

This comprehensive comparison of clade-1 strains suggests the existence of at least four additional novel species, as exemplified by strains 569 (origin unknown), DD13 (from sugarcane), LMG 9002 (from orange), F4 (origin unknown), 3498 (origin unknown), LMG 8992 (from orange), and NCPPB 1128 (from common bean) (Figure 1; Supplemental Tables 1 and 2) (Aritua et al. 2015; Bansal et al. 2020). Notably, for many isolates it is not clear whether they are pathogenic on their host of isolation or on other plants, or if they are regular constituents of the plant’s microbiome, perhaps even with protective features. This genome-wide comparison challenges previous views of the phylogenetic relationships within clade-1 and supports recent findings based on comparing 466 core genes (Rana et al. 2022). Comparison of 16S rRNA sequences grouped *X. sacchari* and ‘*X. sontii’* in one subclade and *X. albilineans, X. hyacinthi, X. theicola*, and *X. translucens* in another subclade (Hauben et al. 1997; Bansal et al. 2021). In contrast, comparisons based on 16S–23S rDNA intergenic spacer sequences or on a portion of the gyrase B gene suggested that *X. albilineans, ‘X. pseudalbilineans’*, *X. sacchari*, and ‘*X. sontii’* form one subclade and *X. hyacinthi, X. theicola*, and *X. translucens* belong to another subclade (Gonçalves and Rosato 2002; Parkinson et al. 2009; Khenfous-Djebari et al. 2019; Koebnik et al. 2021). Our genome-wide ANI comparison clearly separates *X. albilineans* and ‘*X. pseudalbilineans’* from the rest of the clade-1 strains, which correlates with the reduced genome size of these two species (average 3.7 Mbp for 20 strains) in comparison to the other 55 strains in our comparison (4.9 Mbp) (Pieretti et al. 2009; 2015).

## Biosynthesis of coronatine-like compounds in *Xanthomonas*

The phytotoxin coronatine (COR) consists of two moieties, the polyketide coronafacic acid (CFA) and the cyclopropyl amino acid coronamic acid (CMA), which are conjugated by an amide bond. COR and COR-like molecules are produced by several pathovars of *Pseudomonas syringae*, but also some plant-pathogenic enterobacteria and streptomycetes (Bender et al. 1996; Slawiak & Lojkowska 2009; Fyans et al. 2015). In *P. syringae*, COR production is under control of the unconventional two-component system CorRPS, which mediates COR biosynthesis at 18 °C, whereas COR is not detectable at 28 °C (Smirnova et al. 2002). COR and COR-like molecules have been identified as structural mimics of the plant hormone jasmonoyl-L-isoleucine (JA-Ile) (Bender et al. 1999; Bown et al. 2017), which affects plant signaling pathways with importance for plant development and defense (Gimenez-Ibanez et al. 2015). Notably, COR was identified as the virulence factor in *P. syringae* that is responsible for suppressing stomatal innate immune defense (Melotto et al. 2006).

Because *X. campestris* pv. *phormiicola* was reported to synthesize coronatine-like molecules, we searched the genome sequences for homologs of the nine *cfa* genes (coronafacic acid biosynthesis), seven *cma* genes (coronamic acid biosynthesis), the *cfl* gene (coupling factor), and the three regulatory *cor* genes from *Pseudomonas syringae* pv. *tomato* strain DC3000 using TBLASTN. Homologs were detected for all nine Cfa sequences (between 66% and 92% sequence identity), with the Cfl sequence (64% sequence identity), and with the three Cor sequences (between 58% and 67% sequence identity). In contrast, only CmaA and CmaT had hits with 32% sequence identity in both cases, which were, however, not associated with coronamic acid biosynthesis.

We then further looked whether related genes are found in other bacteria, using the CFBP 8444 Cfa, Cfl and Cor amino acid sequences as queries. As expected, we found homologs for the whole biosynthetic gene set in species of *Pseudomonas*, in several enterobacteria (e.g. *Brenneria, Pectobacterium, Dickeya, Lonsdalea*), and in *Streptomyces*. However, the regulatory *cor* genes were only found in *Pseudomonas*. Surprisingly, we also identified close homologs of all these genes in two additional strains of *Xanthomonas*, annotated as *X. arboricola* strain 3058 and *X. campestris* strain 3075. According to the BioProject descriptions at NCBI (PRJNA583332, PRJNA583333), these strains originated from a study of plant-associated saprophytic bacteria and their role in plant health and plant-pathogen interactions. However, since these two *Xanthomonas* strains share a genome-wide ANI of 98.7%, they belong to the same species. Based on analyses at the Type Strain Genome Server (https://tygs.dsmz.de/) (Meier-Kolthoff & Göker 2019), both strains are predicted to belong to a hitherto undescribed species within clade 2 of *Xanthomonas*.

The regulatory *cor* genes were also identified in the genome sequence of *X. theicola*, strain CFBP 4691. However, none of the *cfa* or *cfl* genes were found. Since the Cor regulatory system serves as a thermosensor in *P. syringae* we speculate that biosynthesis of a COR-like substance is likely controlled in a similar way in *Xanthomonas*, but not in enterobacteria or *Streptomyces*, while other factors might be under thermal control in *X. theicola*.

## A novel flagellar gene cluster in *Xanthomonas*

*Xanthomonas* has been proposed as a name for a group of non-sporing, rod-shaped, uni-or rarely biflagellate or non-motile Gram-negative bacteria, forming abundant yellow, slimy colonies on nutrient agar and potato, and mostly digesting starch and producing acid in lactose but not in salicin (Dowson 1939). Genomics has since confirmed the presence of a canonical gene cluster for flagella biosynthesis in almost all sequenced strains, regardless of the species. Yet, some strains have experienced deletion mutations in the flagellar gene cluster which rendered them non-motile on diagnostic media (Darrasse et al. 2013; Jacobs et al. 2015). All three genomes (Table 1) encoded the canonical flagellar gene cluster, which has the same overall genomic organization as the one in clade-2 strains (Supplemental Figure 1).

Surprisingly, strain CFBP 8444 was found to possess a second gene cluster for the biosynthesis of alternative or additional flagella (Figure 3). This gene cluster is unique in the genus *Xanthomonas*. The only other *Xanthomonas* that was found to have a homologous system, strain XNM01, has been misclassified and belongs to the sister genus *Pseudoxanthomonas* (Gowda et al. 2022). BLASTP searches revealed more homologs in additional strains of *Pseudoxanthomonas*, in one strain of *Pseudomonas*, in several beta proteobacteria (mainly from the order Burkholderiales, but also from Neisseriales, Nitrosomonadales and Rhodocyclales) and in one strain of alpha proteobacteria (*Sphingomonas*). This atypical gene cluster contains a flagellin gene, named *fliC* or *lafA*, that is sometimes annotated as lateral flagellin. Overall, sequence similarity between the predicted CFBP 8444 flagellar proteins and their homologs is not very strong with less than 70% sequence identity for the LafA (FliC) protein and on average 64% sequence identity to the most similar system in strain XNM01.

**Figure 3.**
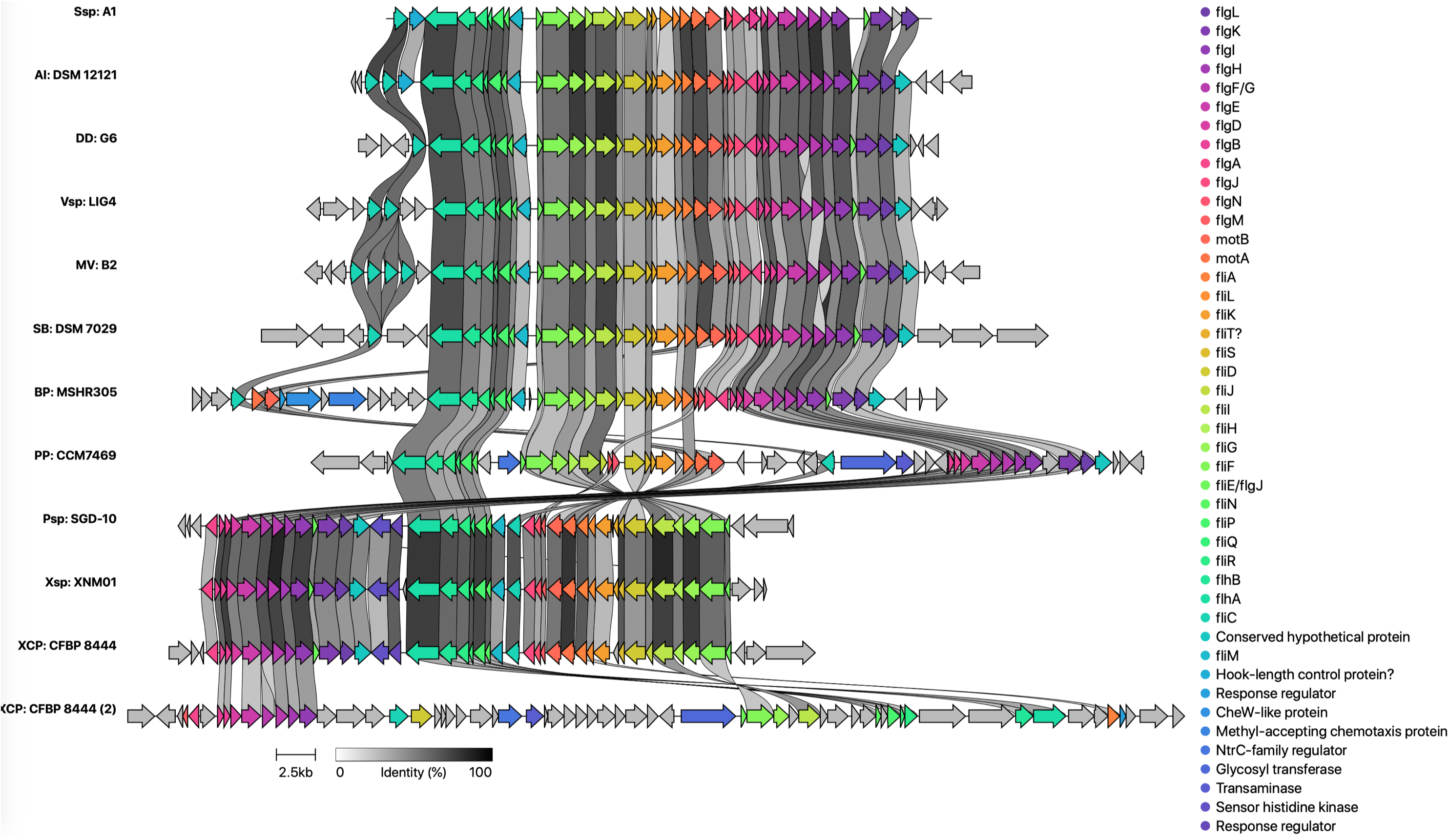
Comparison of flagellar gene clusters. Corresponding regions, extracted from GenBank files, were compared and visualized with clinker (Gilchrist & Chooi 2021). The following strains and genomes were analyzed (from top to bottom): *Sphingomonas* sp. strain A1 (LC043068), *Azoarcus indigens* strain DSM 12121 (SNVV01000001), *Dechloromonas denitrificans* strain G6 (CP075186), *Vogesella* sp. strain LIG4 (LT607802), *Massilia violaceinigra* strain B2 (CP024608), *Schlegelella brevitalea* strain DSM 7029 (CP011371), *Pseudomonas panipatensis* strain CCM 7469 (FNDS01000003), *Pseudoxanthomonas* sp. strain SGD-10 (RRYL01000001), *Xanthomonas* sp. strain XNM01 (JACVAP010000008), and *Xanthomonas campestris* pv. *phormiicola* strain CFBP 8444 (CP102593). Strain CFBP 8444 encodes two flagellar gene clusters, with the novel cluster above the canonical cluster (2), which is shown at the bottom.

*Vibrio parahaemolyticus* is another bacterium that encodes two types of flagella, which are both associated with a distinct cell types, the swimmer cell and the swarmer cell (McCarter & Silverman 1989). The swimmer cell is a short rod that harbors a single polar flagellum, which is produced when the bacterium is grown in liquid media. On semi-solid media, swarmer cells of *V. parahaemolyticus* are elongated and synthesize, in addition to the polar flagellum, numerous lateral flagella which are responsible for translocation over surfaces. Transition between swimmer and swarmer cells is mediated by contact with surfaces and involves the polar flagellum as a tactile sensor. In addition, iron limitation serves as a second signal that is required for swarmer cell differentiation (McCarter & Silverman 1989).

Mesophilic aeromonads are another group of bacteria that express a polar flagellum in all culture conditions, and certain strains produce lateral flagella on semisolid media or on surfaces (Canals et al. 2006). In addition to motility, lateral flagella of *Aeromonas* species have been found to mediate epithelial cell adherence and biofilm formation (Gavín et al. 2002). Later, homologs of the lateral flagellin were found in many beta proteobacteria, such as *Burkholderia dolosa* (Roux et al. 2018). Notably, a *lafA* deletion mutant was more motile under swimming conditions due to an increase in the number of polar flagella, but was not affected in biofilm formation, host cell invasion, or murine lung colonization or persistence over time. *Shewanella putrefaciens* is another species that possesses two separate flagellar systems (Kühn et al. 2022). Using functional fluorescence tagging it was found that screw thread-like motility is mediated by the primary, polar flagellum, and that the lateral flagella support spreading through constricted environments such as polysaccharide matrices.

Lateral-type flagellins were also found in the alpha proteobacterium *Sphingomonas* (Maruyama et al. 2015). *Sphingomonas* sp. strain A1, originally identified as a non-motile and aflagellate bacterium, possesses two sets of flagellar genes, consisting of 35 and 46 genes, respectively. The smaller set encodes the lateral-type flagellin, whereas the larger set includes two flagellar genes typical for polar flagella (Maruyama et al. 2015). Strain A1 cells became motile when they were cultured on semisolid media, due to the presence of a single flagellum at the cell pole, which surprisingly consisted of both lateral and non-lateral flagellins (Maruyama et al. 2015). Immunogold labeling later demonstrated that strain A1 produces two types of flagellar filaments, one formed by three flagellins (lateral and non-lateral flagellins) and the other formed only by lateral flagellins (Kobayashi et al. 2016). Interestingly, lateral-type flagellins were found at the proximal end of the chimeric filaments, next to the hook basal body, whereas the polar flagellins were detected further apart from the cell surface (Kobayashi et al. 2016).

Given the low sequence conservation between the CFBP 8444 noncanonical flagellar genes and those in other proteobacteria, it will be interesting to study this phenomenon further. It remains to be explored whether this strain of *Xanthomonas* produces chimeric polar flagella or lateral flagella under certain conditions, and if so, which environmental clues trigger a switch between different types of flagella.

## Conclusion

The first complete genome sequence for three undescribed species will stimulate further work on the underexplored clade-1 of xanthomonads. Strikingly, this work revealed the presence of a novel, noncanonical flagellar gene cluster that may produce lateral flagella, challenging the view of *Xanthomonas* as a unipolar flagellated bacterium. This work also presents the genetic basis for the production of COR-like molecules in the genus *Xanthomonas*. Since coronatine has been shown to stimulate re-opening of plant leaves’ stomata during the infection process, it will be interesting to study the contribution of these molecules to pathogenicity. Notably, strain CFBP 8444, which synthesizes these COR-like molecules, does not produce type 3 effectors, which in other xanthomonads suppress stomatal immunity (Wang et al. 2021; Liu et al. 2022; Raffeiner et al. 2022).

## Acknowledgements

We are grateful to Cécile Dutrieux and Audrey Lathus for assistance with strains preservation. The authors acknowledge the ISO 9001-certified IRD i-Trop HPC (South Green Platform) at IRD Montpellier for providing high-performance computing resources. Chloé Peduzzi is supported with a FRIA (Formation à la Recherche dans l’Industrie et l’Agriculture) fellowship from the Fonds de la Recherche Scientifique (FNRS). This article is based upon work from COST Action CA16107 EuroXanth, supported by COST (European Cooperation in Science and Technology).”

**e-Xtra Supplemental Table 1. Heatmap of pairwise genome-wide average nucleotide identities (ANI) between clade-1 xanthomonads.** Genome sequences were downloaded from GenBank (accession numbers given in row 3) and pairwise ANI were calculated on a Galaxy-implemented version of FastANI (Jain et al. 2018). This analysis included all publicly available genome sequences from clade 1, except for *X. translucens*, where only eight pathotype strains were chosen as representatives of the genetic diversity of this species (Goettelmann et al. 2022). Names of strains are given in the first column and in the second row. The first row indicates the respective *Xanthomonas* species. SLC, species-level clade (Parkinson et al. 2009).

**e-Xtra Supplemental Table 2. Pairwise genome-wide average nucleotide identities (ANI) and digital DNA-DNA hybridization (dDDH) values of selected clade-1 xanthomonads.**

Pairwise average nucleotide identities (ANI) were calculated on a Galaxy-implemented version of FastANI (Jain et al. 2018). ANI values are given in the upper triangle of the comparison matrix, in blue color. Digital DNA-DNA hybridization values were calculated at the TYGS/GGDC platform (https://ggdc.dsmz.de) (Meier-Kolthoff et al. 2013, 2022). GLM-based DDH estimate, as estimated by the recommended formula 2 (identities / HSP length), are given in the lower triangle of the comparison matrix, in red color. Cells with values that are considered to indicate that the two compared genomes belong to strains of the same species are highlighted in green. Cells with values that are in the ‘grey zone’ of species delineation are in light green. GenBank accession numbers given in row 3.

**e-Xtra Supplemental Figure 1. Comparison of canonical flagellar gene clusters.** Corresponding regions, extracted from GenBank files, were compared and visualized with clinker (Gilchrist & Chooi 2021). The following strains and genomes were analyzed (from top to bottom): *Xanthomonas* sp. strain CFBP 8443 (CP102592), *Xanthomonas campestris* pv. *phormiicola* strain CFBP 8444 (CP102593), *Xanthomonas* sp. strain CFBP 8445 (CP102594), and *Xanthomonas euvesicatoria* pv. *euvesicatoria* strain 85-10 (AM039952).

## Notes

### Competing Interest Statement

The authors have declared no competing interest.

