## Supplementary figures and images for "Novel genetic traits in *Xanthomonas* unveiled by the complete genome sequencing of three clade-1 xanthomonads"

### Supplemental Figure 1

#### Slide 1
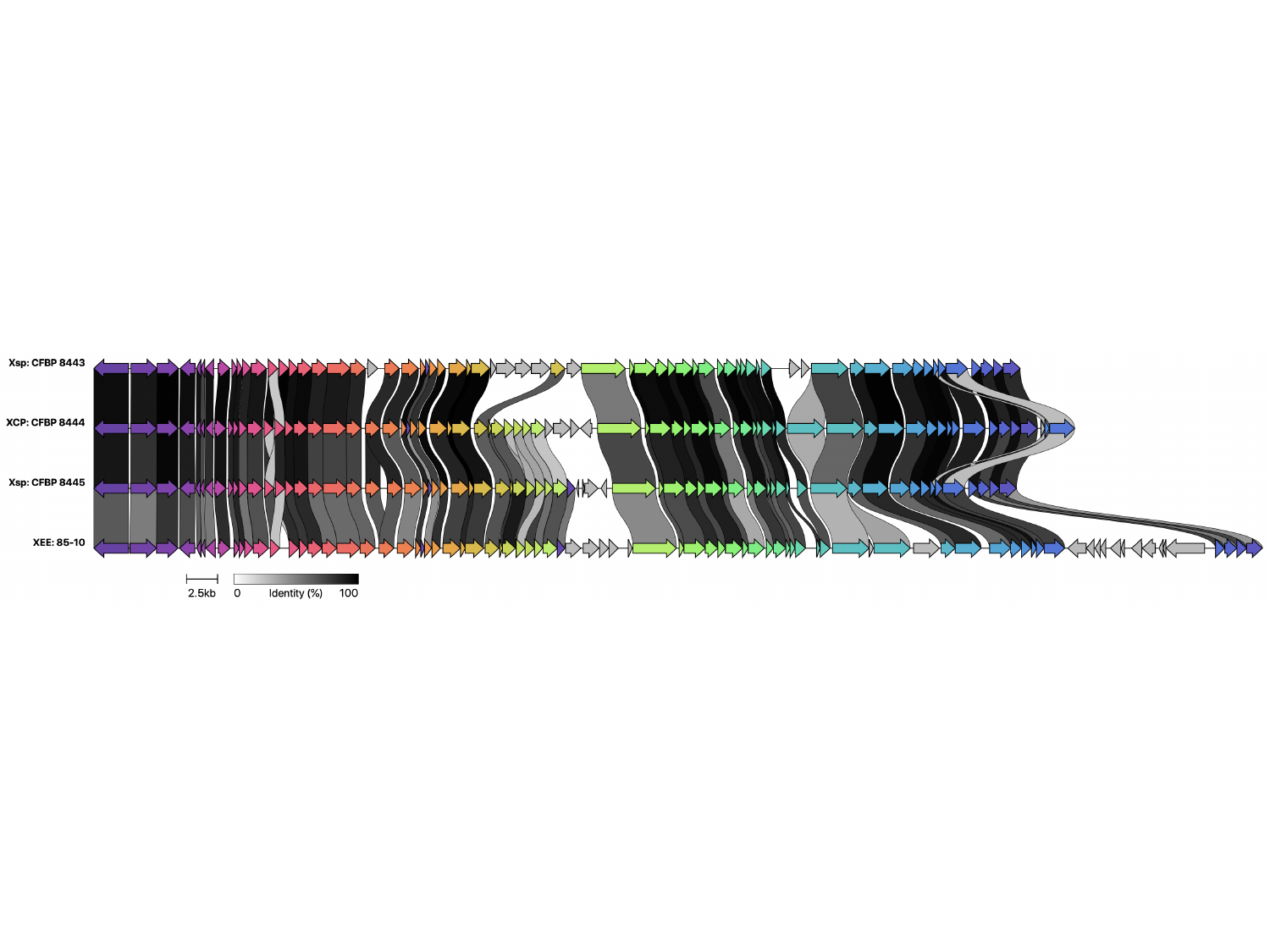
